# A simple theory to explain super-additivity of highly similar drug combinations

**DOI:** 10.1101/2025.10.27.684841

**Authors:** Emily Weed-Nichols, James M. Seckler, Paulina M. Getsy, Stephen J. Lewis, Alan Grossfield

## Abstract

Small-molecule drugs canonically act by binding to a specific site on a single protein, leading to a change in the protein’s activity. In particular, allosteric agonists bind preferentially to the active form of the protein, increasing the population of the active state and increasing activity. Hence, it is not surprising that similar compounds often act in similar ways, because they naturally bind to the same sites. However, recent work has provided examples of closely related small molecules that act super-additively when co-administered, a phenomenon that is difficult to explain using this approach. Here, we derive a simple thermodynamic model that describes how a super-additive response can occur, even for two very similar ligands. We discuss its implications for the specific case of cysteine-derived compounds and the treatment of opioid withdrawal symptoms and suggest avenues by which it could be tested experimentally.

## 1 Introduction

The effects of small molecule drugs are generally determined by their interactions with a target protein or proteins, driven by their specific structure and chemistry. Similar structures tend to lead to similar interactions via binding to the same binding sites, although the resulting effects can differ, e.g. competitive inhibitors are often structurally similar to agonists. It is not uncommon for enantiomers of small molecules to have distinct effects (Lu, 2007; Lapicque et al., 1993; Mehvar et al., 2002). In some cases, the final drug formulations contain a mixture of enantiomers, while others require chiral specificity. The stimulant Adderall, commonly used in the treatment of Attention Deficit Hyperactivity Disorder (ADHD), is an example of both; some components are a single stereoisomer, while others are racemic mixtures, because a non-racemic mixture out-performs both the pure d-isomer and an equal mix of the l- and d-forms(Joyce et al., 2007).

In cases where compositions mix several very similar molecules, such as the enantiomers discussed above, one expects the effects of the various components to be essentially additive, since they are working via the same mechanism. However, there are a few examples in the literature where two enantiomeric molecules act super-additively when co-administered (Borden et al., 1976; Itoh et al., 1997), including recent work on a pair of enantiomeric cysteine derivatives used in the treatment of opioid-induced respiratory depression and withdrawal (Getsy et al., 2022a,b; Baby et al., 2024; Getsy et al., 2025). This kind of super-additivity is difficult to explain using the standard single protein-ligand interaction model of drug action.

In this paper, we develop a simple thermodynamic model to explain this phenomenon, based on the binding polynomial formulation of Wyman and Gill (Wyman and Gill, 1990). The central idea is that two distinct pathways activated by the small molecules; each molecule is a high-affinity agonist for one and a low-affinity agonist for the other, and both pathways must be active to see a physiological response. We formally derive the model, demonstrate its behavior under different conditions, and discuss the assumptions and limitations of the model as presently formulated. Finally, we consider the implications for the specific cases of cysteine derivatives used for the treatment of opioid withdrawal symptoms (Getsy et al., 2025), and suggest avenues for experimental verification.

## 2 Model for super-additivity

### 2.1 Basic theory of allostery

We will derive the basic theory of allostery using the binding polynomial formalism of Wyman and Gill (Wyman and Gill, 1990). Let *P* be a protein, which can exist in active or inactive states, denoted *P*_*A*_ and *P*_*I*_. The equilibrium between these states is given by the equation

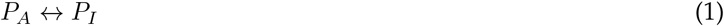

controlled by the equilibrium constant, *K*_eq_ = [*P*_*A*_]*/*[*P*_*I*_]. For a ligand *L* that only binds to the active state, we have a second equation

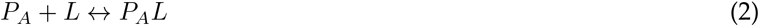

where the binding constant is defined by

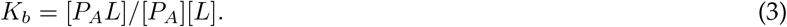

If we assume that the active forms of the protein with and without ligand are indistinguishable, which is expected for allosteric ligands, we can write the apparent equilibrium constant as

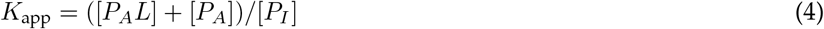

If we rewrite this in terms of the *K*_*b*_ by substituting for [*P*_*A*_*L*] using Equation 3, we get

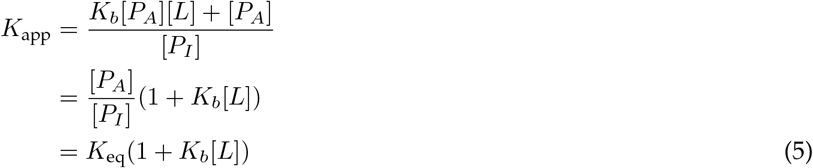

To get the probability of an active conformation

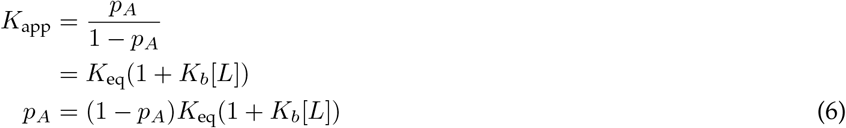

Solving for *p*_*A*_ yields

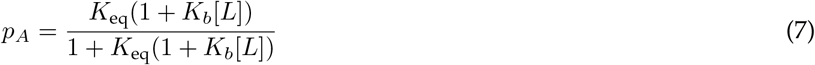

This demonstrates the well-known result that the probability of the active state increases with increasing agonist concentration, shown in Figure 1. In the absence of ligand, the protein exists almost entirely in the inactive state, but as the ligand concentration increases, so too does the protein’s activity. Ligand 1 takes effect at a lower concentration, as expected since we chose for it to have a binding constant 100x smaller than Ligand 2.

**Figure 1:**
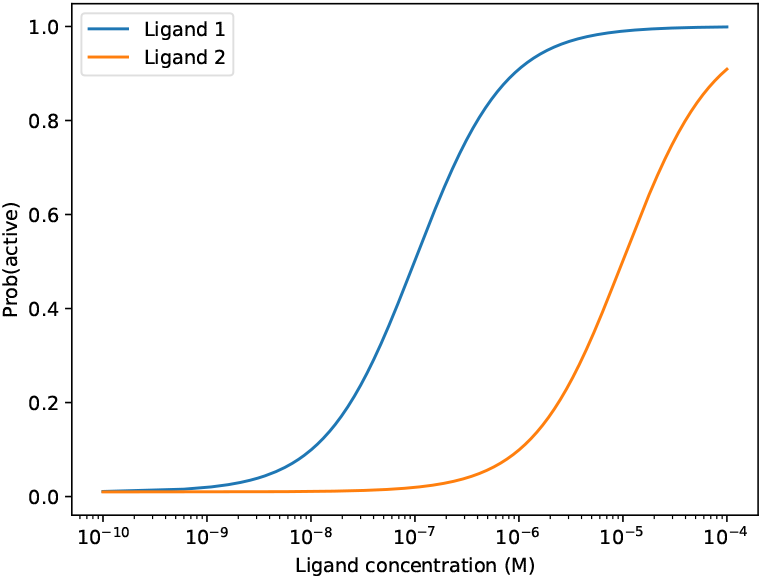
The probability of the active state of a protein as a function of ligand concentration, calculated using Equation 7. The protein equilibrium constant was set to 0.01, while *K*_*b*_ = 1 nM for Ligand 1 and *K*_*b*_ = 100 nM for Ligand 2.

The whole derivation can be trivially generalized to the case of multiple ligands *L*_1_ and *L*_2_ with binding constants *K*_*b*1_ and *K*_*b*2_ to yield

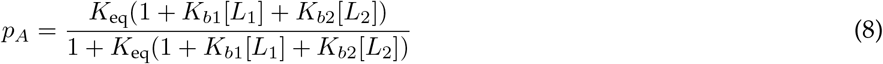

where *K*_*b*1_ and *K*_*b*2_ are the binding constants for Ligands 1 and 2, respectively. In the limit where Ligands 1 and 2 have similar binding coefficients, the effects of the two ligands are additive.

### 2.2 Model for super-additivity

The above derivation does not in itself explain how a pair of highly similar compounds could produce super-additive effects. However, it does suggest a mechanism by which this could occur: if the physiological effects of the ligands are the result of two distinct signaling pathways, both of which must be activated to produce the response, the probability of activating each of the individual pathways can be computed using Equation 8. If we further assume the two pathways function independently, the overall fractional response will be the product of the two individual probabilities, computed as

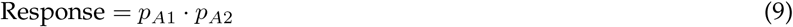

As a concrete example, if Ligand 1 is a strong agonist for the first pathway but only a weak agonist for the second, then physiologically it will appear to be a weak agonist: at low ligand concentration, pathway 1 will be turned on, but the physiological response will be absent because pathway 2 is not activated, since we assume both pathways must be active to produce a response. If Ligand 2 is a weak agonist for the first pathway but a strong agonist for the second, the same phenomenon will be observed, because while pathway 2 will be activated, pathway 1 will not.

Apparent super-additivity emerges when both ligands are present. According to Equation 8, if either Ligand 1 or Ligand 2 binds tightly, that will be sufficient to activate the corresponding pathway. In the case described above, where each ligand is a strong agonist for one pathway and a weak agonist for the other, the result will be that both pathways are activated at low ligand concentrations, leading to a super-additive physiological response.

This is neatly demonstrated by Figure 2. Panels 2A and 2B show the probability of activating the first and second proteins, respectively, as a function of the concentration of the two ligands. The difference in the ligand’s binding constants is readily apparent as the “white rectangle” indicating the region where the protein is not active. Panel 2C shows the overall response, computed using Equation 8, essentially the product of the two previous plots. Proceeding from zero concentration of both ligands (the lower left corner of the plot), we see that one can obtain a response at much lower concentration by proceeding along the diagonal (increasing the concentration of both ligands simultaneously) than by increasing the concentration of either ligand alone.

**Figure 2:**
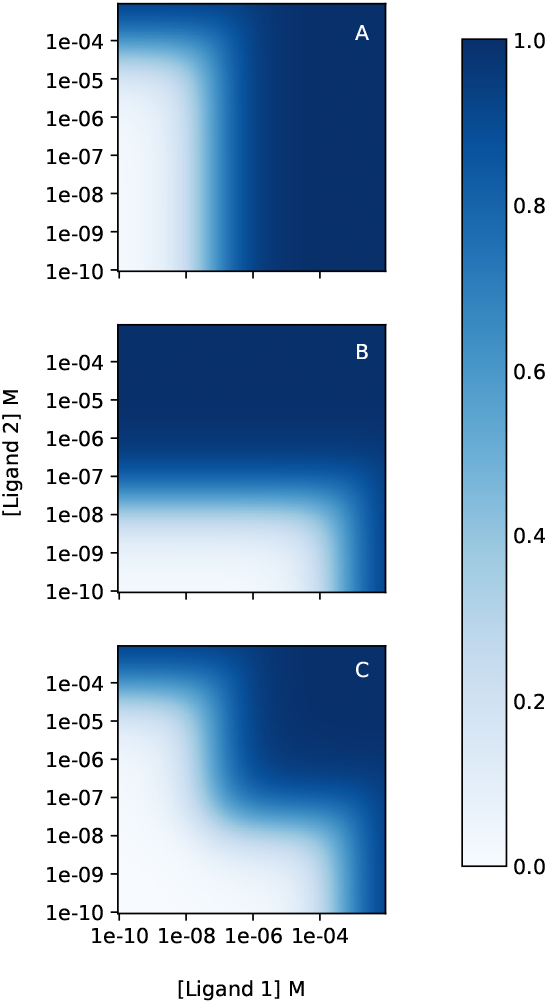
Overall signal response as a function of ligand concentration. Panel A shows the probability for the first protein to be active, computed using Equation 8. Panel B shows the same, for the second protein. Panel C shows the overall functional response, computed using Equation 9. The protein equilibrium constant was set to 0.01 for both proteins. For the first protein, *K*_*b*_ was set to 0.1 nM for Ligand 1 and *K*_*b*_ = 100 nM for Ligand 2. For the second protein, the values were reversed. The intensity of the color indicates the probability that one or both pathways will be active, our proxy for the amplitude of the signal response.

The results are simpler to view if we reduce the plots to a single dimension. Figure 3 shows the same data as Figure 2, but only considering 2 cases. In Figure 3A, we show the response of the individual proteins and the overall response to Ligand 1. These plots are equivalent to tracing along the X-axis of Figure 2, with the concentration of Ligand 2 set to zero. The equivalent plot for Ligand 2 would look identical, but with the Protein 1 and Protein 2 curves exchanged. Figure 3B shows the same quantities, but for the case where the concentrations of Ligand 1 and Ligand 2 are equal. This is equivalent to walking along the diagonal of Figure 2. In this case, the super-additive effects of combining the two ligands are readily apparent: in the case where a single ligand is present, the response reaches 50% at [Ligand] ≈ 10^−5^M, while when both ligands are present, the equivalent response occurs at roughly 100x lower ligand concentration.

**Figure 3:**
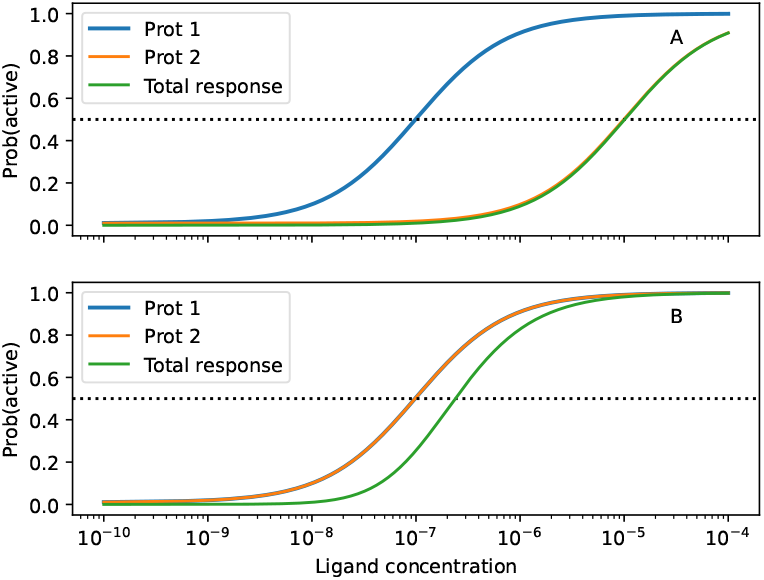
Overall signal response as a function of ligand concentration. Panel A shows 3 curves: the response of the first protein to Ligand 1, the response of the second protein to Ligand 1, and the overall response to Ligand 1; the total response curve is almost superposable with the response of protein 2, since at concentrations where protein 2 is active, protein 1 is almost entirely in the active state. Panel B shows the equivalent curves where the concentration of Ligand 1 and Ligand 2 are equal (the x-axis value is their sum). In each plot, there is a horizontal dotted line to mark 50% activity or response.

The full model is depicted pictorially in Figure 4. For those who are curious to see the effects of the specific choices for binding constants used above, we have made available an interactive Jupyter notebook at https://colab.research.google.com/drive/1ltd4Bkz9lgAFP2lQ8DaFV4KcUlv6iBtC. A short version of the derivation is presented, with plots similar to the ones presented here. However, the notebook figures contain sliders that allow the user to interactively change the binding constants and immediately see the effects on the activity and overall response.

**Figure 4:**
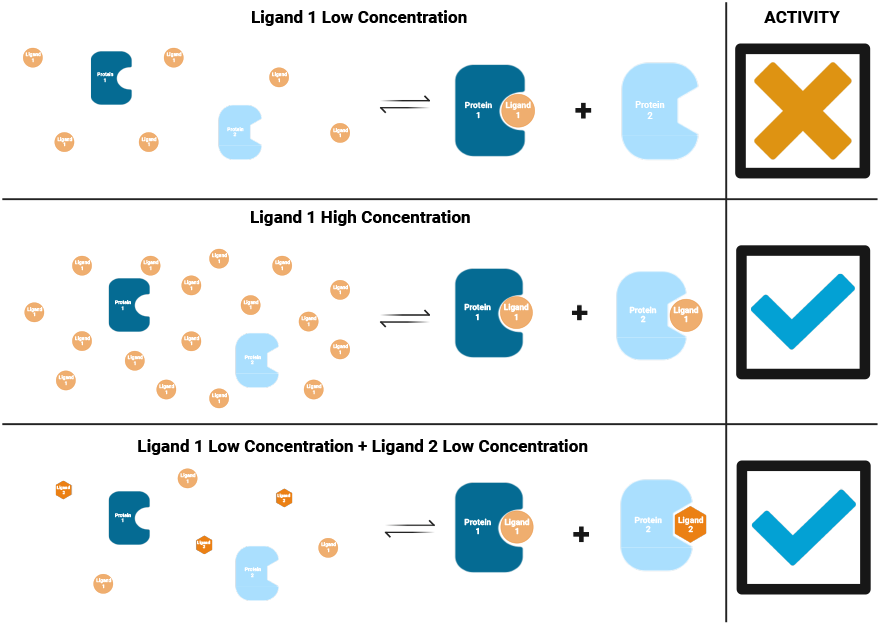
Pictorial depiction of the model for super-additive function. Created in BioRender. Weed-Nichols, E. (2025) https://BioRender.com/0fyhmqh

## 3 Discussion

### 3.1 Connection between microscopic *K*_*b*_ and EC_50_

There is a tendency in the literature and elsewhere to equate the EC_50_, the concentration at which a drug has half its maximal effect, with its microscopic *K*_*b*_ or binding constant. While this is an intuitive notion, Figure 1 shows that it is not in fact correct. For instance, Ligand 1 has a *K*_*b*_ of 1 nM, while the probability of the protein being in the active state doesn’t reach 0.5 until the ligand concentration is 100 nM. The explanation is revealed by Equation 7: in addition to the binding constant and the ligand concentration, the protein’s intrinsic equilibrium constant also plays a role. The lower the protein equilibrium constant — the less favorable the active state is in the absence of the ligand — the more ligand is required to stabilize the active state. For the purposes of Figure 1, we chose *K*_eq_ = 0.01, a sufficiently low value that activity would be undetectable by most assays in the absence of the allosteric ligand. A larger value would shift the curves in Figure 1 to the left, while smaller values would shift them to the right. Put another way, because in this model we’ve chosen to assume the ligand only binds to the active state, the overall free energy of binding is the sum of the free energy required to convert the protein to the active state and the free energy gained when the ligand binds to it. The *K*_*b*_ in Equation 3 is the microscopic binding constant, the affinity of the ligand to the active state, so it only reports on the latter component of the free energy. This is how *K*_*b*_ would be computed using molecular simulation techniques, such as docking (Pagadala et al., 2017) or alchemical ligand binding calculations (Mey et al., 2020) based on the active state. That said, methods that take the full protein equilibrium into account do exist (Smith et al., 2024). However, this is not what would be measured in a functional EC_50_ assay, where the readout is presumed to be proportional to the population of the active state, or even in a binding assay, where the readout is the total amount of ligand bound.

### 3.2 Model limitations and opportunities for generalization

The model described here is intended as a simple model to illustrate a potential mechanism by which two similar compounds could have super-additive effects. As such, it is clearly a simplification, and numerous details and potential alternatives are left out.

For example, the derivation is written assuming that the ligands function as allosteric agonists that turn on a signaling pathway. However, the model functions equally well if one or both ligands are inverse agonists, which bind to the inactive state to turn off an already active pathway; this could be a pathway activated by a challenge, or something constitutively active. The equivalence can be seen trivially by reversing the roles of *P*_*A*_ and *P*_*I*_ in Equations 3 to 7. The resulting derivation produces essentially the same expressions, except that now increasing ligand concentration will decrease the probability of the active state rather than increasing it.

A somewhat stronger assumption made in the above derivation was that the ligands bind have no affinity for the protein when it is in the inactive state, and only bind when it is active. It is not difficult to generalize the derivation to remove this assumption, and the resulting equation for the probability of the active state (the equivalent of Equation 7) is not that much more complicated. However, Equation 8 already has 4 binding constants as free parameters in addition to *K*_eq_, and the general form would have another 4. The equation for super-additivity, Equation 9, has 10 free parameters (one equilibrium constant and four binding constants for each protein), so the general form would have 18. It’s not clear whether one could uniquely fit dose-response data using the simpler model derived here, so the additional complexity does not seem worthwhile. The only real advantage to generalizing the derivation is that it would simultaneously treat both agonists and inverse agonists. However, this does not produce any new physical insights, beyond the notion that two very similar ligands can have additive, super-additive, or competitive effects depending on their respective affinities for the active and inactive states. Since additive agonism and competitive antagonism are already well-studied phenomena, and the active-state binding model is sufficient to describe the origins of super-additivity, we focus here on the simpler model.

Finally, Equation 9 is presented as the behavior when two distinct proteins that bind the ligands are involved, each activating a different pathway. That said, the model would also apply if there were multiple binding allosteric sites on a single protein that could activate multiple pathways. *β*-arrestin is one such protein, which activates numerous pathways and is differentially trafficked after inducing the internalization of an activated GPCR (Gurevich, 2014).

In this case, the choice to simply multiply the probabilities of the two individual active states — essentially assuming that the two active states are totally independent of each other — would become less justified, since it’s likely that there would be some coupling between the two activities, even if they involve distinct binding sites and active sites. Moreover, depending on the trafficking required for activity, the two activities might compete against each other. One could include this in the derivation via a coupling term between the two binding sites — essentially introducing the concept of cooperative binding — but there would be no novel insights and would require yet another free parameter.

### 3.3 Applications of this model to cysteine esters and opioid-induced respiratory depression

The model presented here makes the most sense for the case where the two ligands are chemically very similar; we expect that similar ligands will bind to the same site on a protein, although we expect different affinities and possibly different effects (e.g. one could be an agonist and the other an inverse agonist or neutral antagonist). If the two compounds are not similar, it would be difficult to justify the assumption that they bind to the same site on each of two different proteins. This limits the general applicability of the model and shows that it cannot explain all cases where two drugs have super-additive physiological effects.

However, one example where the model makes sense is the case of the d- and l-enantiomers of cysteine ethyl ester (CYSee) or related compounds. Both compounds counter opioid-induced respiratory depression in rats (Getsy et al., 2022a; Baby et al., 2024), with comparable efficacies. Recent work showed that if the enantiomers are given together, they exert their clinical effects at 5-10x lower doses than either compound alone (Getsy et al., 2025). The two compounds have very similar effects physiologically, and given their structural similarity, shown in Figure 5, it is reasonable to assume they act via the same mechanism. Thus, it is difficult to explain their super-additive effects while invoking only a single protein binding site, while the model proposed here provides a plausible framework.

**Figure 5:**
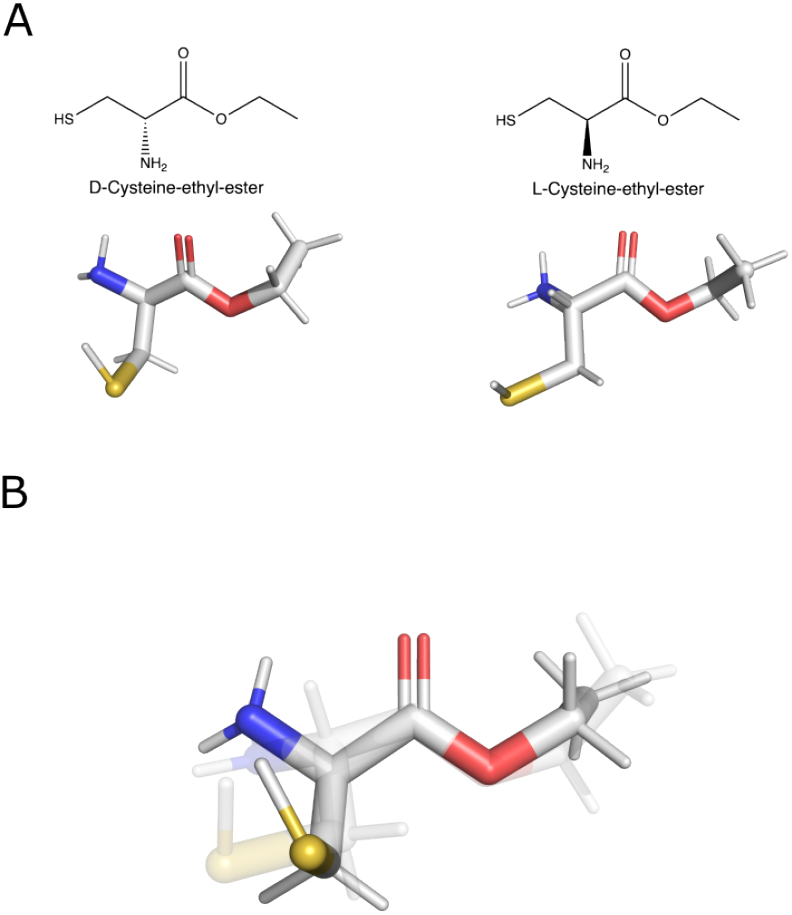
Panel A shows the chemical structures of d- and l-CYSee. Panel B shows the two structures superimposed, to emphasize their similarity.

In the context of the present model, we propose that d-CYSee and l-CYSee each act via 2 distinct pathways; each is a high-affinity agonist for one response and a low-affinity agonist for the other. For example, we are currently exploring the hypothesis that they act by binding to *β*-arrestin(data not shown), altering the trafficking of the *µ*-opioid receptor after activation and thus changing downstream signaling. As part of its normal function, *β*-arrestin must form multiple protein-protein interactions, first with clathrins to facilitate GPCR internalization, and later with other proteins to direct *β*-arrestin between recycling and degradation, and turn on various downstream signaling partners (Gurevich, 2014). It is possible that there are multiple allosteric binding sites on *β*-arrestin that separately modulate these interactions. Alternatively, binding could occur on a separate protein, such as HINT1 or the NMDA receptor, that also participates in the response to opioids (Rodríguez-Muñoz et al., 2010).

Testing this hypothesis in the context of whole-cell experiments is difficult. A more direct (if still challenging) approach would be to measure the affinities between the two ligands and a purified protein of interest (e.g. *β*-arrestin2) via a technique such as isothermal titration calorimetry or surface plasmon resonance.

## 4 Conclusions

In this manuscript, we derive a model to explain how two highly similar compounds can have super-additive effects when co-administered, based on the notion that they bind to two distinct allosteric sites, both of which must be activated to produce a physiological response. Although in principle this model could be used to fit dose-response data, in all likelihood its primary value will be to suggest alternative experiments to search for a secondary target.

